# Spatiotemporal dynamics of memory encoding and memory retrieval states

**DOI:** 10.1101/2022.06.15.496281

**Authors:** Yuju Hong, Isabelle L. Moore, Devyn E. Smith, Nicole M. Long

## Abstract

Memory encoding and memory retrieval are neurally distinct brain states that can be differentiated on the basis of cortical network activity. However, it is unclear whether sustained engagement of one network or fluctuations between multiple networks give rise to these memory states. The spatiotemporal dynamics of memory states may have important implications for memory behavior and cognition; however, measuring temporally resolved signals of cortical networks poses a challenge. Here we recorded scalp electroencephalography from subjects performing a mnemonic state task in which they were biased toward memory encoding or retrieval. We performed a microstate analysis in order to measure the temporal dynamics of cortical networks throughout this mnemonic state task. We find that microstate E, a putative analog of the default mode network, shows temporally sustained dissociations between memory encoding and retrieval, with greater engagement during retrieve compared to encode trials. We further show that decreased engagement of microstate E is a general property of encoding, rather than a reflection of retrieval suppression. Together, these findings show that memory states are supported by sustained engagement of a particular cortical network. Memory success, as well as cognition more broadly, may be influenced by the ability to engage or disengage microstate E in a goal-dependent manner.

## Introduction

Successful memory depends on both the formation of a representation of a new experience – encoding – and subsequently accessing the stored representation of a past experience – retrieval. Although brain activity patterns during encoding are commonly reinstated during retrieval (Wheeler, Petersen, & Buckner, 2000; Polyn, Natu, Cohen, & Norman, 2005; Ritchey, Wing, LaBar, & Cabeza, 2013), there is growing evidence that memory encoding and retrieval constitute distinct brain states (Hasselmo, 2005; Richter, Chanales, & Kuhl, 2016) supported by cortical network activity (Long & Kuhl, 2021). However, it is unclear whether memory states are supported by the sustained engagement of a specific network or rapid fluctuations between distinct networks. Memory states influence processing and behavior (Long & Kuhl, 2019, 2021; Smith, Moore, & Long, 2022); encoding and retrieval failures may be due to over or under reliance on one or more cortical networks. The aim of this study is to investigate the spatiotemporal dynamics of memory states.

Attention, control, memory, and other cognitive demands recruit distinct resting state networks (RSNs). Identified via resting state functional magnetic resonance imaging (fMRI), RSNs reflect nodes of the cortex whose activity is intrinsically correlated (Fox et al., 2005; Power et al., 2011; Yeo et al., 2011). RSNs show both sustained and transient activity fluctuations in response to task demands. The ventral attention (VAN) and frontoparietal control (FPCN) networks are engaged at different moments in time during decision making tasks (Gratton et al., 2016) as are the default mode network (DMN) and the dorsal attention network (DAN) during arithmetic tasks (Raccah, Daitch, Kucyi, & Parvizi, 2018). The DMN, commonly ‘deactivated’ during cognitive tasks (Raichle et al., 2001), is engaged during episodic memory retrieval (Kim, 2010), semantic processing (Binder, Desai, Graves, & Conant, 2009), and internally directed attention (Buckner & DiNicola, 2019). There is likewise evidence that communication between RSNs supports cognitive processing, in particular that connectivity between the FPCN and the DMN supports internally directed attention (Kam et al., 2019), and other forms of internal mentation, including working memory (Murphy, Bertolero, Papadopoulos, Lydon-Staley, & Bassett, 2020) and memory search (Kragel & Polyn, 2013). Together, these findings broadly point to a role for RSNs in cognition, but leave open the question of how the temporal dynamics of RSNs support memory states.

Encoding and retrieval states can be distinguished via multivariate pattern classification analysis of cortical signals. Although there exist multiple definitions of “brain state” (“unique configuration of anatomy, physiology, representation and behavior,” Kay & Frank, 2019; “recurring patterns of correlation between networks,” Beaty et al., 2018; “the reliable patterns of brain activity that involve the activation and/or connectivity of multiple large-scale brain networks,” Tang, Rothbart, & Posner, 2012), the general theme across these definitions is that brain *states* are inherently multivariate in nature. That is, a single brain region, neuronal population, and/or electrical signal is unlikely to be the sole substrate responsible for a particular brain state. Consistent with the interpretation that encoding and retrieval reflect brain *states*, we have found that encoding and retrieval recruit distinct whole-brain activity patterns. Spectral activity across electrodes placed on the scalp (Long & Kuhl, 2019; Smith et al., 2022) as well as fMRI BOLD activity across cortical regions and networks (Richter et al., 2016; Long & Kuhl, 2021) differentiate encoding from retrieval states. However, these findings do not elucidate whether memory states are supported by sustained or transient engagement of RSNs. To address this gap, it is necessary to adopt an analytical approach that yields time-resolved signals from cortical networks.

Temporal dynamics of RSNs can be measured through the investigation of *microstates* (Lehmann, Ozaki, & Pal, 1987). Microstates are global patterns of voltage scalp topographies that remain stable for a sustained period of time (approximately 60 to 120 ms; Michel & Koenig, 2018). Four canonical microstates (A, B, C, D) have been identified across both resting and task-based studies, although there is evidence for additional microstates (E, F, G and sub-divisions of C) beyond the canonical four. Studies employing simultaneous EEG and fMRI recordings, as well as source-localization methods, indicate that EEG microstates are closely associated with fMRI RSNs, including the visual, attention, and default mode networks (Britz, Van De Ville, & Michel, 2010; Custo et al., 2017; Michel & Koenig, 2018; Yuan, Zotev, Phillips, Drevets, & Bodurka, 2012). For instance, Microstate B shows increased engagement during eyes-open compared to eyes-closed rest (Seitzman et al., 2017) and is linked with the visual network (Britz et al., 2010). Microstate D, thought to reflect the DAN (Britz et al., 2010), is engaged during a serial subtraction task (Seitzman et al., 2017) and processes that (re)orient attention (Milz et al., 2016). Finally, both Microstate C and E have been linked with the DMN (Bréchet et al., 2019; Custo et al., 2017) and show increased engagement during eyes-closed rest, memory retrieval and mind wandering (Koenig et al., 2002; Bréchet et al., 2019). Together, microstates provide a method for assessing the temporal dynamics of cortical networks.

Our hypothesis is that memory brain states are supported by distinct patterns of cortical topography or microstates that are sustained over time. The alternative is that fluctuations between multiple different microstates give rise to encoding and retrieval states. To test our hypothesis, we recorded scalp EEG while subjects engaged in a mnemonic state task (Smith et al., 2022). Briefly, subjects were instructed to either encode the currently presented stimulus or to retrieve a previously presented, related stimulus. These instructions were intended to bias subjects toward either an encoding or retrieval state. Using established methods (Murray, Brunet, & Michel, 2008), we measured microstate engagement over time as subjects were biased toward these memory states.

## Materials and Methods

### Subjects

109 (65 female, age range = 18-66, mean age = 21.64 years), fluent English speakers from the University of Virginia community participated. All subjects had normal or corrected-to-normal vision. Informed consent was obtained in accordance with the University of Virginia Institutional Review Board for Social and Behavioral Research and subjects were compensated for their participation. Nine subjects were excluded from the final dataset: one had previously participated in a behavioral version of the experiment, one had poor EEG signal quality, two failed to comply with task instructions, two had corrupt data files, and three had no resting state data. Thus, the reported data is from the remaining 100 subjects. A subset of this dataset was previously reported (Smith et al., 2022); all of the analyses and results described here are novel. The raw, de-identified data and the associated experimental and analysis codes used in this study will be made available via the Long Term Memory laboratory website upon publication.

### Mnemonic State Task Experimental Design

Stimuli consisted of 576 object pictures, drawn from an image database with multiple exemplars per object category (Konkle, Brady, Alvarez, & Oliva, 2010). From this database, we chose between 108 to 216 unique object categories and two exemplars from each category. For each subject, one exemplar served as a List 1 object, one as a List 2 object; object condition assignment was randomly generated for each subject.

#### General Overview

Subjects completed between 6 to 12 runs in which they viewed two lists containing 18 object images. For the first list, each object was new (List 1 objects). For the second list (List 2 objects), each object was again new, but was categorically related to an object from the first list. For example, if List 1 contained an image of a bench, List 2 would contain an image of a different bench (Figure 1). During List 1, subjects were instructed to encode each new object. During List 2, each trial contained an instruction to either encode the current object (e.g. the new bench) or to retrieve the corresponding object from List 1 (the old bench). Following the study phase, subjects completed a recognition phase which tested subjects’ memory of List 1 and List 2 objects. Because subsets of our subjects completed different types of recognition memory tests, we do not report the memory test data here.

**Figure 1.**
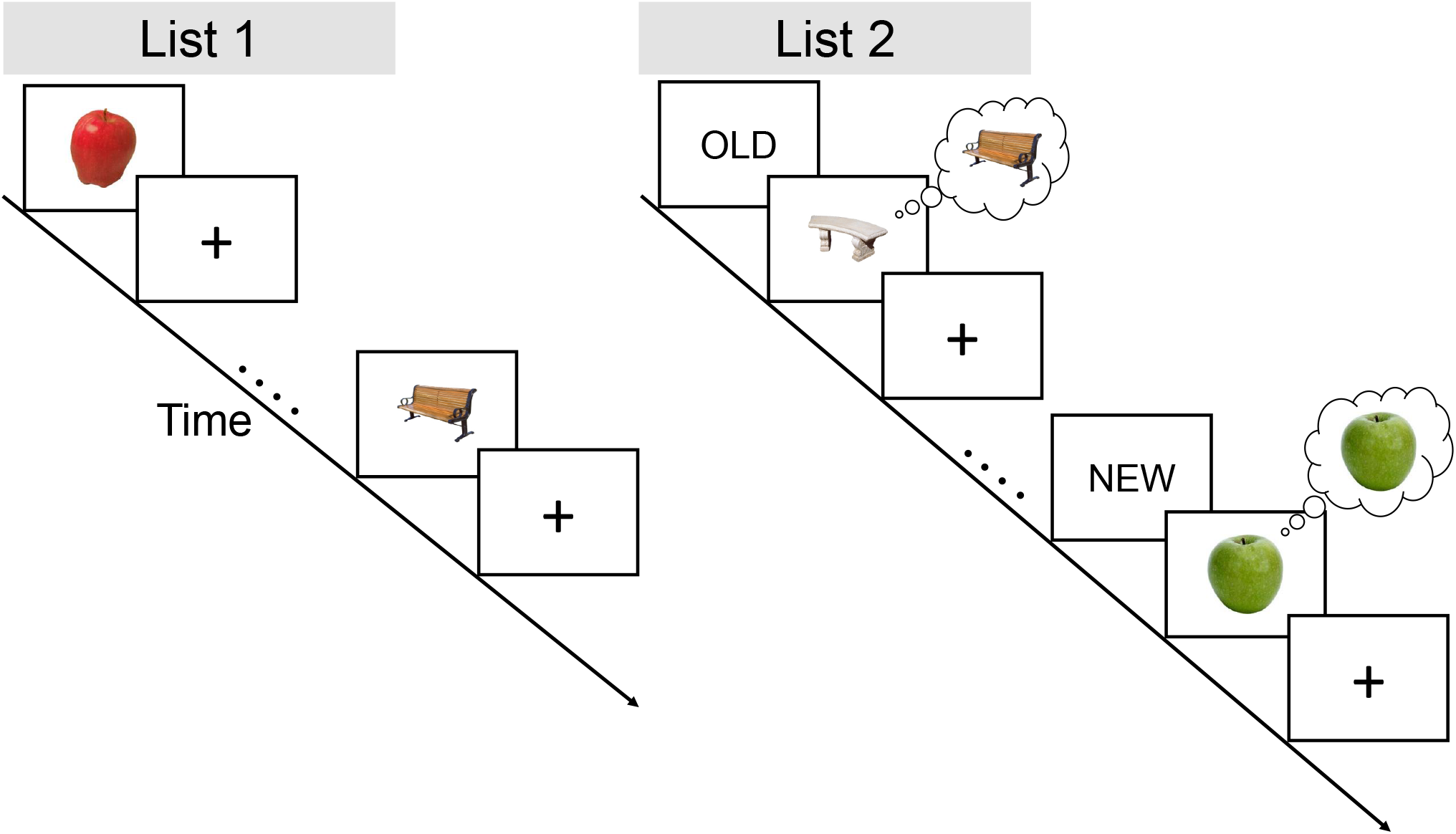
Mnemonic State Task Design. During List 1, subjects studied individual objects (e.g. bench, apple) for 2000 ms followed by a 1000 ms inter-stimulus interval. During List 2, subjects saw novel objects that were from the same categories as the objects shown in List 1 (e.g., a new bench, a new apple). Preceding each List 2 object was an “OLD” instruction cue or “NEW” instruction cue. The “OLD” cue signaled that subjects were to *retrieve* the corresponding object from List 1 (e.g., the old bench). The “NEW” cue signaled that subjects were to *encode* the current object (e.g. the new apple). Each run of the experiment contained a List 1 and List 2; object categories (e.g., bench) were not repeated across runs.

#### List 1

On each trial, subjects saw a single object presented for 2000 ms followed by a 1000 ms inter-stimulus interval (ISI). Subjects were instructed to study the presented object in anticipation of a later memory test.

#### List 2

On each trial, subjects saw a cue word, either “OLD” or “NEW” for 2000 ms. The cue was followed by presentation of an object for 2000 ms, which was followed by a 1000 ms ISI. All objects in List 2 were non-identical exemplars drawn from the same category as the objects presented in the immediately preceding List 1 of the same run. That is, if a subject saw a bench and an apple during List 1, a different bench and a different apple would be presented during List 2. On trials with a NEW instruction (encode trials), subjects were to encode the presented object. On trials with an OLD instruction (retrieve trials), subjects tried to retrieve the categorically related object from the preceding List 1. Importantly, this design prevented subjects from completely ignoring List 2 objects following OLD instructions in that they could only identify the to-be-retrieved object category by processing the List 2 object.

Subjects completed 6 to 12 runs with two lists in each run (List 1, List 2). Subjects viewed 18 objects per list, yielding a total of 216 to 288 object stimuli from 108 to 216 unique object categories. Subjects did not make a behavioral response during either List 1 or List 2.

### Resting State Task

After the mnemonic state task, the subjects completed a simple and cognitively non-demanding resting state task (Figure 2). The task consisted of 8 runs with 10 trials per run. During each trial, subjects viewed five asterisks presented on a computer screen, with one asterisk at the center of the screen and the four others at the top, bottom, left, and right sides of the screen relative to the center. Subjects performed one of four types of eye movements during each run, with the specific type indicated by an auditory cue. During open/close runs, subjects were instructed to open or close their eyes. During left/right runs, subjects were instructed to saccade to the left or right asterisk. During up/down runs, subjects were instructed to saccade to the top or bottom asterisk. During the blink runs, subjects were instructed to blink following the auditory cue. All auditory cues within a run were separated by an ISI. For 41 subjects, the ISI was a fixed interval of 5000 ms. For 59 subjects, the ISI could vary randomly between 3000 ms and 9000 ms with an average of 6500 ms. For all runs except blink runs, the auditory cues alternated (e.g. “open,” “close,” “open,” “close”) and were evenly divided into five of each subtype.

**Figure 2.**
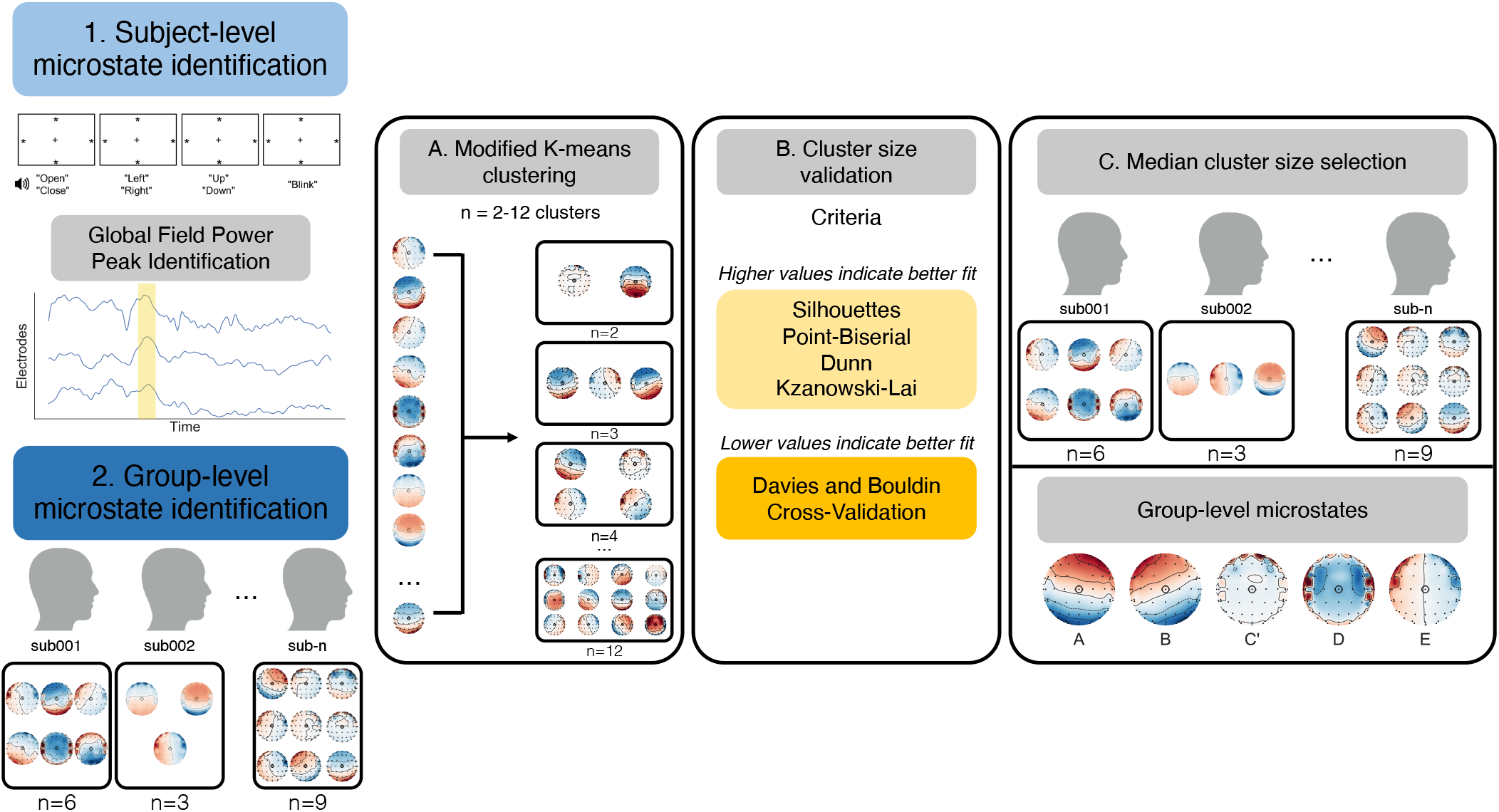
Microstates: Procedure and Results. **(1)** We first identified subject-level microstates. From each subject’s resting state data, we extracted the top 1000 peaks of global field power. **(A)** We submitted these peaks to a modified *k* -means clustering algorithm with the number of possible clusters ranging from 2 to 12. **(B)** We used six criteria, Silhouettes, Davies and Bouldin, Point-Biserial, Dunn, Kzanowski-Lai (for *n* clusters > 2), and Cross Validation, to evaluate the fit of these 2 to 12 clusters. Each criterion provided a best fitting number of clusters. For the criteria in yellow (Silhouettes, Point-Biserial, Dunn, Kzanoski-Lai), higher values indicate better fit; for the criteria in orange (Davies and Bouldin, Cross-Validation), lower values indicate better fit. **(C)** Cluster size selection was performed by selecting the median best fitting *n* clusters across the six criteria. As an example, if the six criteria resulted in the best fitting *n* clusters: n = 2, 5, 6, 8, 10, 12, the median cluster size selection would be n = 7. The outcome of this step resulted in a single *n* cluster set that best fit the subject’s resting state data. **(2)** After identifying the best *n* fitting microstates for each subject, we subjected the subject-level microstate maps to the same procedure of **(A)** modified *k* -means clustering followed by **(B)** cluster size validation and **(C)** cluster size selection. As a result of this process, we identified five group-level microstates in the resting state data. We use the naming convention employed in previous research to facilitate interpretation and comparison across datasets.

### EEG Data Acquisition and Preprocessing

EEG recordings were collected using a BrainVision system and an ActiCap equipped with 64 Ag/AgCl active electrodes positioned according to the extended 10-20 system. All electrodes were digitized at a sampling rate of 1000 Hz and were referenced to electrode FCz. Offline, electrodes were later converted to an average reference. Impedances of all electrodes were kept below 50 kΩ. Electrodes that demonstrated high impedance or poor contact with the scalp were excluded from the average reference. Bad electrodes were determined by voltage thresholding (see below).

Custom Python codes were used to process the EEG data. We applied a high pass filter at 0.1 Hz, followed by a notch filter at 60 Hz and harmonics of 60 Hz to each subject’s raw EEG data. We then performed three preprocessing steps (Nolan, Whelan, & Reilly, 2010) to identify electrodes with severe artifacts. First, we calculated the mean correlation between each electrode and all other electrodes as electrodes should be moderately correlated with other electrodes due to volume conduction. We z-scored these means across electrodes and rejected electrodes with z-scores less than -3. Second, we calculated the variance for each electrode as electrodes with very high or low variance across a session are likely dominated by noise or have poor contact with the scalp. We then z-scored variance across electrodes and rejected electrodes with a |z| ≥ 3. Finally, we expect many electrical signals to be autocorrelated, but signals generated by the brain versus noise are likely to have different forms of autocorrelation. Therefore, we calculated the Hurst exponent, a measure of long-range autocorrelation, for each electrode and rejected electrodes with a |z| ≥ 3. Electrodes marked as bad by this procedure were excluded from the average re-reference. We then calculated the average voltage across all remaining electrodes at each time sample and re-referenced the data by subtracting the average voltage from the filtered EEG data. We used wavelet-enhanced independent component analysis (Castellanos & Makarov, 2006) to remove artifacts from eyeblinks and saccades.

### EEG Data Microstate Analysis

We applied a bandpass filter of 1 to 40 Hz and then downsampled the artifact-corrected mnemonic state task and resting state data to 250 Hz. Our microstate analysis, performed via custom Python code based on a publicly available microstate toolbox (Poulsen, Pedroni, Langer, & Hansen, preprint), consists of four main steps (Figure 2). First, we identified subject-level microstates from each subject’s resting state data. Second, we identified group-level microstates by aggregating the individual subject microstates. Third, we applied the group-level microstates to each subjects’ mnemonic state data in order to identify sample-specific microstates during memory encoding and memory retrieval. Finally, we extracted microstate properties from the mnemonic state data.

#### Subject-level microstate identification

For each subject’s resting state data, we identified *n*-clusters of microstates using a modified *k* -means clustering algorithm (Murray et al., 2008). First, we calculated the global field power (GFP) and extracted the top 1000 peaks from the GFP. Next, these peaks were submitted to a *k* -means clustering algorithm. We repeated these steps for sets of clusters, with the number of clusters ranging from 2-12 (Zanesco, Denkova, & Jha, 2021). To identify the number of clusters *n* that best fit a subject’s data, we used six criteria: Silhouettes, Davies and Bouldin, Point-Biserial, Dunn, Kzanowski-Lai (for *n* clusters > 2), and Cross Validation criterion. Each of the six criteria provides a fit value for each cluster set and a cluster set is considered the best if it has either the highest value (Silhouettes, Point-Biserial, Dunn, and Kzanowski-Lai) or the lowest value (Davies and Bouldin, and Cross Validation). To identify the best fitting cluster set across the six criteria, we calculated the median of the identified cluster set sizes. Theoretically, different cluster sets could be the best fit for each individual criteria, e.g. *n* = 2, 5, 6, 8, 10, 12; in this example, the best fit would be *n* = 7 clusters. We repeated this process for each subject to identify subject-level microstates.

#### Group-level microstate identification

We next identified group-level microstates. We submitted all subject-level microstate maps to the modified *k* -means clustering algorithm above and identified the best fitting microstates across subjects using the same process as subject-level microstate identification with the same six criteria.

#### Mnemonic state task microstate identification

Next, we identified microstates in the mnemonic state task. First, for each individual subject, we segmented the mnemonic state task trials. The segmentation window began at stimulus onset and spanned the duration of the stimulus (2000 ms), yielding 500 time samples per trial. We then individually correlated each of the 500 time samples in a given trial with each of the group-level microstate maps. As the critical feature for defining a microstate is the relative difference across the scalp and not the direction of the difference, we do not consider the polarity of the electrodes (Michel & Koenig, 2018). We calculated the absolute value of the resulting Pearson’s *r*, and a sample was labeled with the most correlated microstate except in cases in which none of the resulting *r* values exceeded a threshold of 0.5 (Custo et al., 2017). We performed an additional round of thresholding based on the temporal extent of a given microstate. By definition, a microstate is sustained, meaning that the same microstate should be present for multiple consecutive samples. We required a given microstate to be present for six consecutive samples (a minimum of 24 ms; Zanesco et al., 2021). In all other cases (i.e. no *r* values exceeding 0.5 and insufficient contiguous microstates), the sample was unlabelled and excluded from the analysis.

#### Mnemonic state task microstate properties

We analyzed two microstate properties when assessing the extent to which microstates vary across memory brain states: duration and coverage. Duration represents how long, on average, a microstate persists in milliseconds following its onset. Coverage represents the percentage of the time interval of interest that the microstate is active. In the context of our mnemonic state task, if a microstate were active for a total of 1000 ms throughout the 2000 ms trial interval, the coverage of the microstate would be 50%. We aggregated the duration and coverage values across all subjects separately for encode and retrieve trials.

### Statistical Analyses

To assess microstate dissociations across encode and retrieve trials, as well as over time, we conducted repeated measures ANOVAs. We conducted post-hoc ANOVAs and *t* -tests for significant main effects and interactions. For post-hoc comparisons across microstates and across time, we used false discovery rate (FDR; *p* = 0.05) correction (Benjamini & Hochberg, 1995) to correct for multiple comparisons.

## Results

### Five microstates fit the resting state data

Our first goal was to identify microstates in the resting state data. As the resting state task should not induce memory encoding or retrieval and is distinct from the mnemonic state task, these data are ideal for independently identifying microstates. To this end, we identified the best fitting *n* microstates that characterized the group-level resting state data, resulting in the identification of five microstates (Figure 2). To facilitate interpretation and comparison with prior work, we have adopted the naming conventions used in previous research and label these microstates A, B, C’, D, and E (Michel & Koenig, 2018). The five microstates we identified have previously been associated with particular resting state networks (RSNs) identified via source-localization and fMRI data: the auditory network (Microstate A), the visual network (Microstate B), the saliency network (Microstate C’), the dorsal attention network (Microstate D) and the default mode network (Microstate E; Britz et al., 2010; Custo et al., 2017).

### Longer microstate E duration for retrieve compared to encode trials

Our primary goal was to assess the extent to which the five microstates identified in the resting state data dissociate memory encoding and memory retrieval. A longer microstate duration for a specific memory state would suggest that sustained network engagement supports either encoding or retrieval. We measured the duration of each microstate during the 2000 ms List 2 stimulus interval. We find significant dissociations in microstate duration across encode and retrieve trials (Figure 3A). We conducted a 2 × 5 repeated measures ANOVA with instruction (encode, retrieve) and microstate (A, B, C’, D, E) as factors. We excluded 38 subjects who did not have any trials labeled with either microstate C’ (N = 37) or microstate D (N = 1) from the ANOVA. We found a main effect of instruction (*F* _1,61_ = 10.44, *p* = 0.0020, 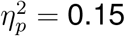). This effect was driven by greater duration across all microstates for retrieve compared to encode trials. We found a main effect of microstate (*F* _4,244_ = 22.79, *p* < 0.001, 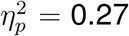) and an interaction between instruction and microstate (*F* _4,244_ = 6.66, *p* < 0.001, 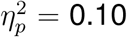). We performed follow-up post-hoc paired *t* -tests to investigate encode vs. retrieve dissociations for each microstate. Microstate duration was significantly greater for retrieve compared to encode trials for microstate E (*t*_61_ = 3.03, *p* = 0.0036, Cohen’s *d* = 0.31; FDR corrected). We found no significant differences in duration for microstates A, B, C’, or D (|*t*’s| < 2.34, *p*’s > 0.027).

**Figure 3.**
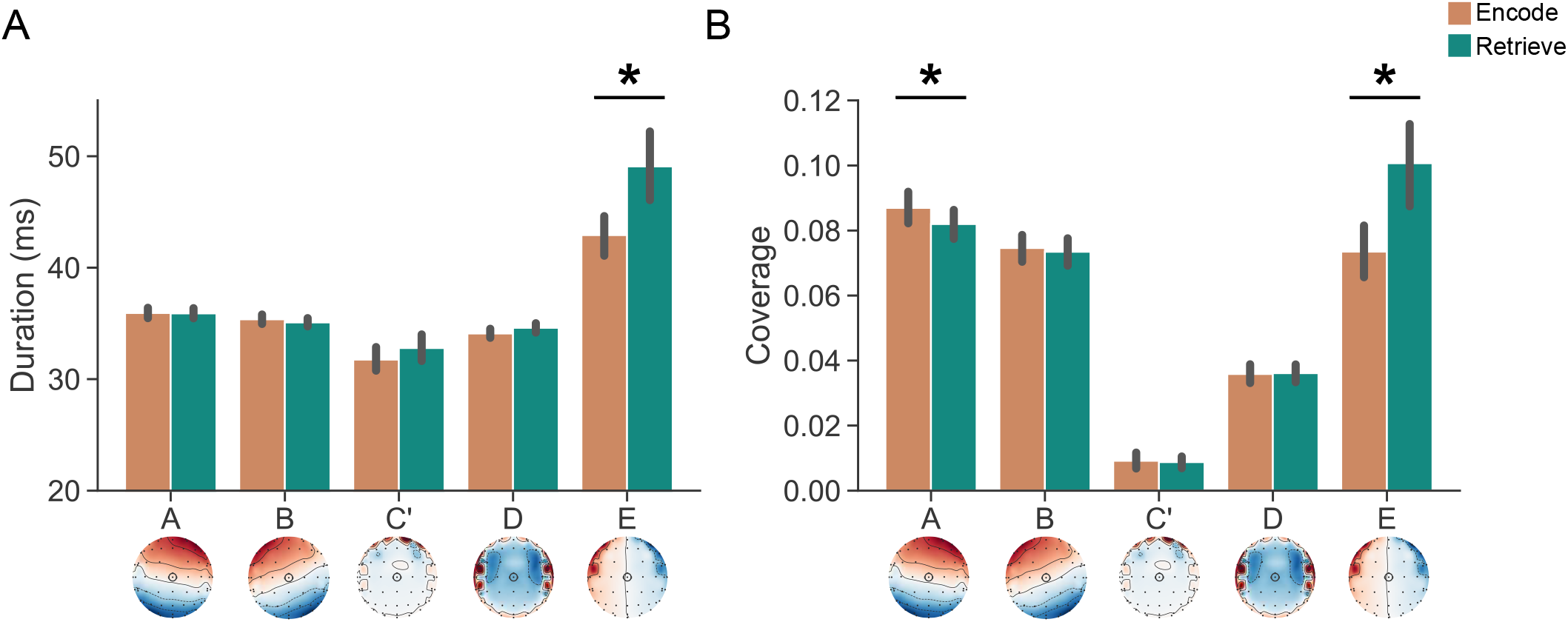
Microstate Duration and Coverage as a Function of Instruction. **(A)** Microstate E has a significantly longer duration during retrieve (teal) compared to encode (orange) trials. **(B)** There is greater microstate A coverage for encode compared to retrieve trials and greater microstate E coverage for retrieve compared to encode trials. Error bars represent standard error of the mean. * denotes the microstates for which the post-hoc *t* -test comparison between encode and retrieve trials survived FDR correction.

We next sought to test whether the increased microstate duration for retrieve trials was specific to microstate E. The interaction between instruction and microstate suggests that duration differences are driven by some, but not all, of the microstates. We computed duration difference scores (retrieve - encode) for each microstate. We found that the duration difference in microstate E (*M* = 6.16, *SD* = 15.90) is significantly greater than all other microstate duration differences (*t*’s > 2.19, *p*’s < 0.033; see Table 1). Together, these results show that the duration of microstate E specifically dissociates encode from retrieve trials.

**Table 1.**
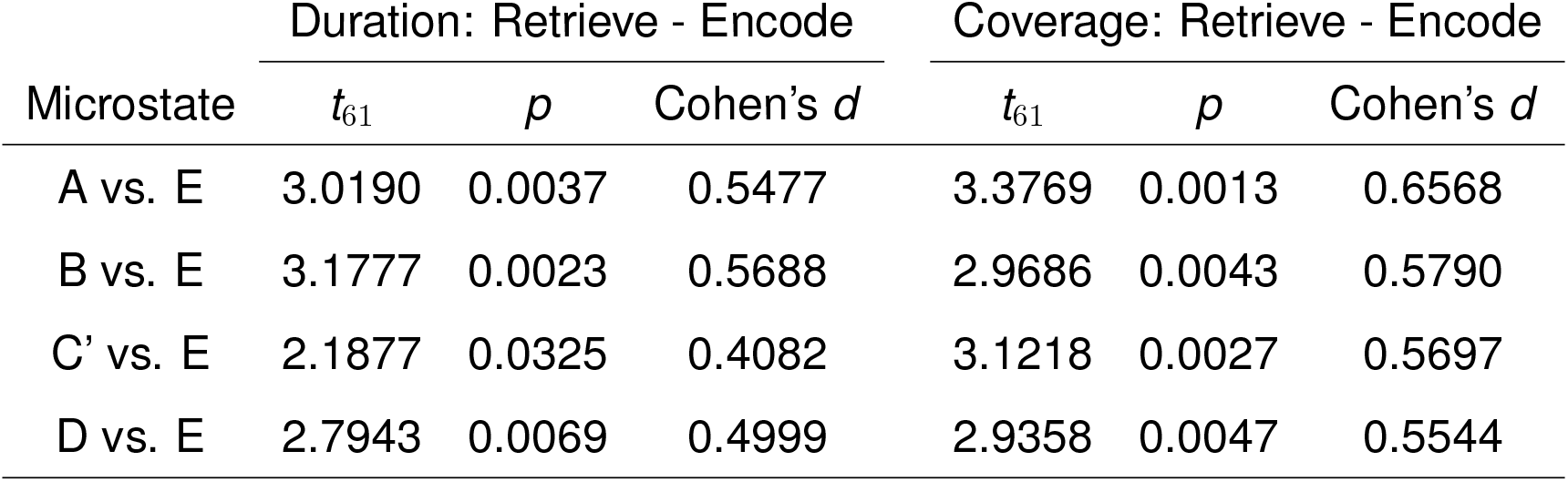
Comparison of Microstate E to Microstates A, B, C’, D.

### Greater microstate E coverage for retrieve compared to encode trials

Having shown that microstate duration varies as a function of instruction, we next sought to assess whether microstate coverage likewise differs across encode and retrieve trials. Greater coverage selectively for encoding or retrieval would suggest that a microstate is more engaged or active during that specific memory state and could be observed even in the absence of a between-condition difference in duration. We measured the coverage of each microstate during the 2000 ms List 2 stimulus interval and found significant dissociations in coverage across encode and retrieve trials (Figure 3B). We conducted a 2 × 5 repeated measures ANOVA with instruction (encode, retrieve) and microstate (A, B, C’, D, E) as factors and coverage as the dependent variable. We found a main effect of instruction (*F* _1,61_ = 8.51, *p* = 0.0049, 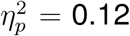), driven by greater coverage across all microstates for retrieve compared to encode trials. We found a main effect of microstate (*F* _4,244_ = 43.38, *p* < 0.001, 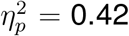) and an interaction between instruction and microstate (*F* _4,244_ = 9.46, *p* < 0.001, 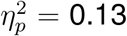). We performed follow-up post-hoc paired *t* -tests to investigate encode vs. retrieve dissociations for each microstate. Microstate coverage is significantly greater for retrieve compared to encode trials for microstate E (*t*_61_ = 3.11, *p* = 0.0028, Cohen’s *d* = 0.32; FDR corrected) and for encode compared to retrieve trials for microstate A (*t*_61_ = 3.34, *p* = 0.0014, Cohen’s *d* = 0.16; FDR corrected). We found no significant differences in coverage for microstates B, C’, or D (*t*’s < 0.80, *p*’s > 0.43).

We next sought to test whether the increased microstate coverage for retrieve trials was specific to microstate E. We computed coverage difference scores (retrieve - encode) for each microstate. We found that the coverage difference in microstate E (*M* = 0.027, *SD* = 0.068) is significantly greater than all other microstate coverage differences (*t*’s > 2.94, *p*’s < 0.0047; see Table 1). Together, these results show that the coverage of microstate E specifically dissociates encode from retrieve trials.

### Sustained engagement of microstate E across retrieve trials

Our analyses above provide strong evidence that encode and retrieve trials are distinguished specifically by the engagement of microstate E. Microstate E is more engaged, both in terms of duration and coverage, during retrieve compared to encode trials. Critically, however, these results do not indicate *when* during the trial microstate E engagement dissociates the two instructions. Microstate E engagement may either be sustained throughout the stimulus interval or its engagement may fluctuate over time. To test these two alternatives, we re-calculated coverage specifically for microstate E in 100 ms time intervals spanning the stimulus interval (2000 ms). Our approach was identical to that used in our prior coverage analysis (see *Methods*), however, we calculated the coverage separately for each of 20 time intervals.

We find a greater dissociation in microstate E coverage between retrieve and encode trials throughout the stimulus interval (Figure 4). We conducted a 2 × 20 repeated measures ANOVA with instruction (encode, retrieve) and time interval as factors and coverage as the dependent variable. We find a main effect of instruction (*F* _1,61_ = 9.66, *p* = 0.0029 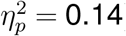), consistent with our coverage analysis above, and a main effect of time interval (*F* _19,1159_ = 19.03, *p* < 0.001, 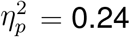). We find an interaction between instruction and time interval (*F* _19,1159_ = 2.89, *p* < 0.001, 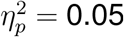). We performed post-hoc paired *t* -tests at each time interval and used FDR correction (*p* = 0.05) to correct for multiple comparisons. The coverage dissociation between encode and retrieve trials was significant at all time intervals excluding the first two hundred milliseconds (0-100 ms, 100-200 ms; |*t*’s| < 1.74, *p*’s > 0.087). Together, these results suggest that memory states are supported by sustained engagement (or disengagement, in the case of encode trials) of microstate E.

**Figure 4.**
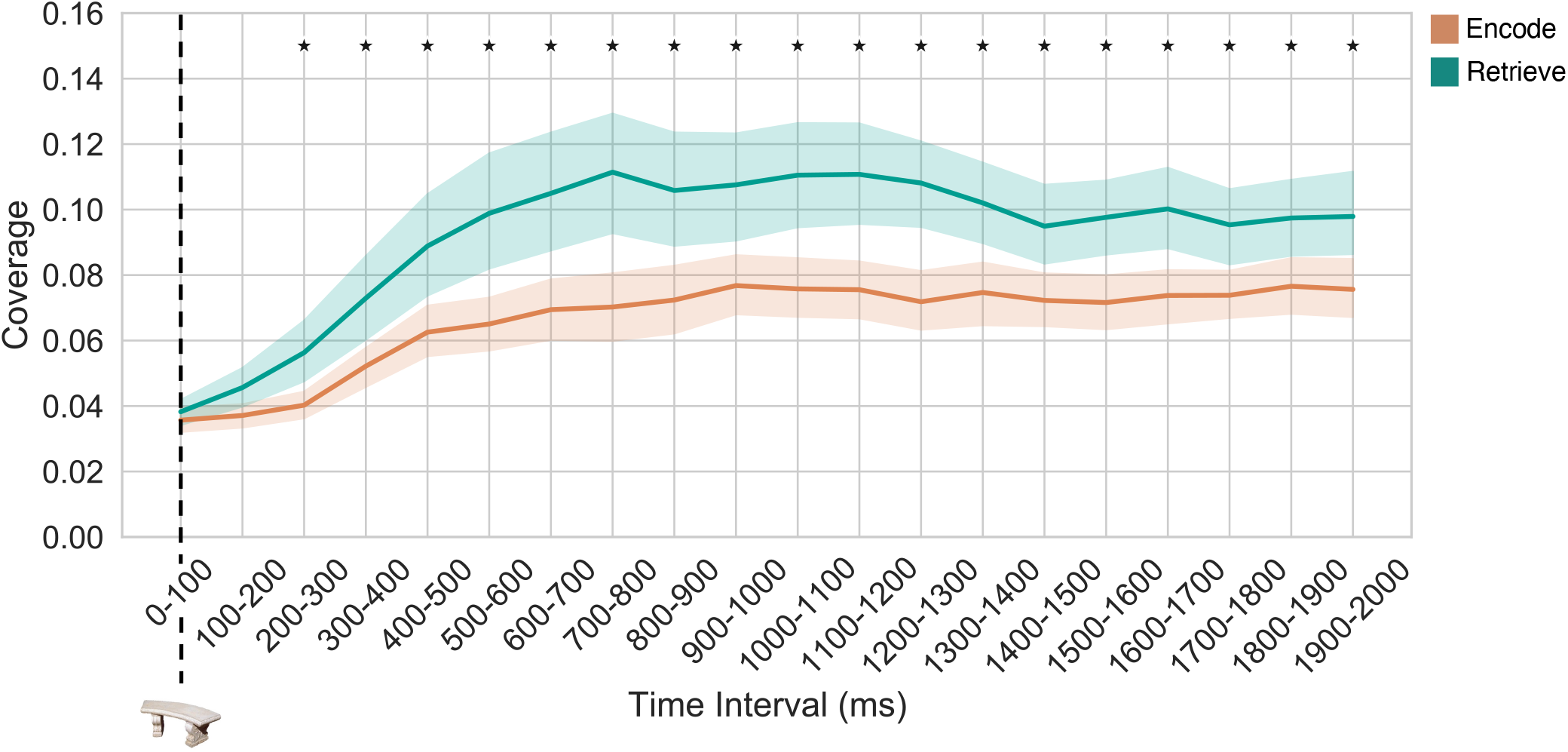
Temporal Dynamics of Microstate E Coverage. We examined microstate E coverage throughout the stimulus interval separately for encode (orange) and retrieve (teal) trials. Stimulus onset at 0-100 ms is indicated by the vertical dashed line. Microstate E coverage is greater during retrieve compared to encode trials for all intervals excluding the first 200 ms. The shaded area represents the standard error of the mean. * denotes time intervals for which the post-hoc *t* -test comparison between encode and retrieve trials survived FDR correction.

### Microstate dissociations between List 1 and List 2 encode trials

Our interpretation is that, compared to retrieval, encoding is characterized by decreased engagement of microstate E. However, it is conceivable that decreased microstate E coverage and duration on encode trials reflects a ‘suppression’ mechanism or a top-down driven inhibition of retrieval, rather than encoding per se. On all List 2 trials, regardless of instruction, there is the potential for retrieval given the presence of recently presented categorically associated stimuli. Indeed, we have found evidence for automatic retrieval of List 1 objects during List 2 trials regardless of instruction (Smith et al., 2022). Therefore, to claim that decreased engagement of microstate E is a general property of memory encoding, it is necessary to measure microstate E engagement when there is little (or no) potential for retrieval. The List 1 trials present just such an opportunity. Although there are no trial-level cues, the List 1 trials are analogous to the List 2 encode trials in terms of memory demands, as subjects were instructed to form a memory of each List 1 object; however, there should be no need for retrieval inhibition on these trials.

To assess the extent to which List 2 encode trials reflect general ‘encoding’ as opposed to interference-driven retrieval-suppression, we measured coverage for all five microstates in 100 ms time intervals spanning the stimulus interval, separately for List 1 and List 2 encode trials (Figure 5). To test for microstate dissociations, we conducted five – one per microstate – 2 × 20 repeated measures ANOVAs with condition (List 1, List 2 encode) and time interval as factors and coverage as the dependent variable (see Table 2). All microstates show main effects of time interval (*F* ‘s > 4.39, *p*’s < 0.001), but only microstate D shows a main effect of instruction (*F* _1,61_ = 4.98, *p* = 0.030, 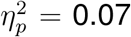) that is driven by greater coverage for List 2 encode trials compared to List 1 trials. Microstates A, D, and E display significant interactions between instruction and time interval (microstate A: *F* _19,1159_ = 1.99, *p* = 0.0071, 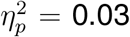; microstate D: *F* _19,1159_ = 3.97, *p* < 0.001, 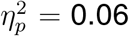; microstate E: *F* _19,1159_ = 1.87, *p* = 0.013, 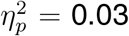). We performed post-hoc paired *t* -tests at each time interval and used FDR correction (*p* = 0.05) to correct for multiple comparisons for each microstate. Microstate A coverage is greater during the 200-300 ms interval on List 1 compared to List 2 trials (*t*_61_ = 3.31, *p* = 0.0016, Cohen’s *d* = 0.33). Microstate D coverage is greater during the 500-700ms time interval on List 2 compared to List 1 trials (500-600ms: *t*_61_ = −3.77, *p* = 0.0004, Cohen’s *d* = 0.45; 600-700ms: *t*_61_ = −3.52, *p* = 0.0008, Cohen’s *d* = 0.37). Microstate E coverage is greater during the 1800-2000 ms interval on List 2 compared to List 1 trials (1800-1900ms: *t*_61_ = −3.02, *p* = 0.0037, Cohen’s *d* = 0.19; 1900-2000ms: *t*_61_ = −3.34, *p* = 0.0014, Cohen’s *d* = 0.17). Although we find some evidence for dissociations in microstate E late in the stimulus interval, microstate E engagement is generally similar across List 1 and List 2 trials (*t*_61_ = 1.07, *p* = 0.290, Cohen’s *d* = 0.05; Bayes Factor = 0.24), suggesting that the List 2 microstate E effects are not merely a suppression of retrieval. List 1 and List 2 encode trials most strongly dissociate in terms of microstate D, which is more engaged, particularly early (500 ms) in the stimulus interval, during List 2 compared to List 1 trials.

**Table 2.** Comparison of microstate coverage over time across List 1 and List 2 encode trials.

| Microstate | Instruction<br>df = (1,61) |  |  | Time Interval<br>df = (19,1159) |  |  | Instruction x Time Interval<br>df = (19,1159) |  |  |
| --- | --- | --- | --- | --- | --- | --- | --- | --- | --- |
| | $F$ value | $p$ value | $\eta_p^2$ value | $F$ value | $p$ | $\eta_p^2$ value | $F$ value | $p$ | $\eta_p^2$ value |
| A | 0.014 | 0.91 | <0.001 | 14.0 | <0.001 | 0.19 | 1.99 | 0.007 | 0.03 |
| B | 0.249 | 0.62 | <0.001 | 16.6 | <0.001 | 0.21 | 1.38 | 0.125 | 0.02 |
| C' | 0.074 | 0.79 | 0.001 | 4.39 | <0.001 | 0.07 | 1.47 | 0.089 | 0.02 |
| D | 4.98 | 0.03 | 0.07 | 5.36 | <0.001 | 0.08 | 3.97 | <0.001 | 0.06 |
| E | 1.14 | 0.29 | 0.02 | 26.6 | <0.001 | 0.30 | 1.87 | 0.013 | 0.03 |

**Figure 5.**
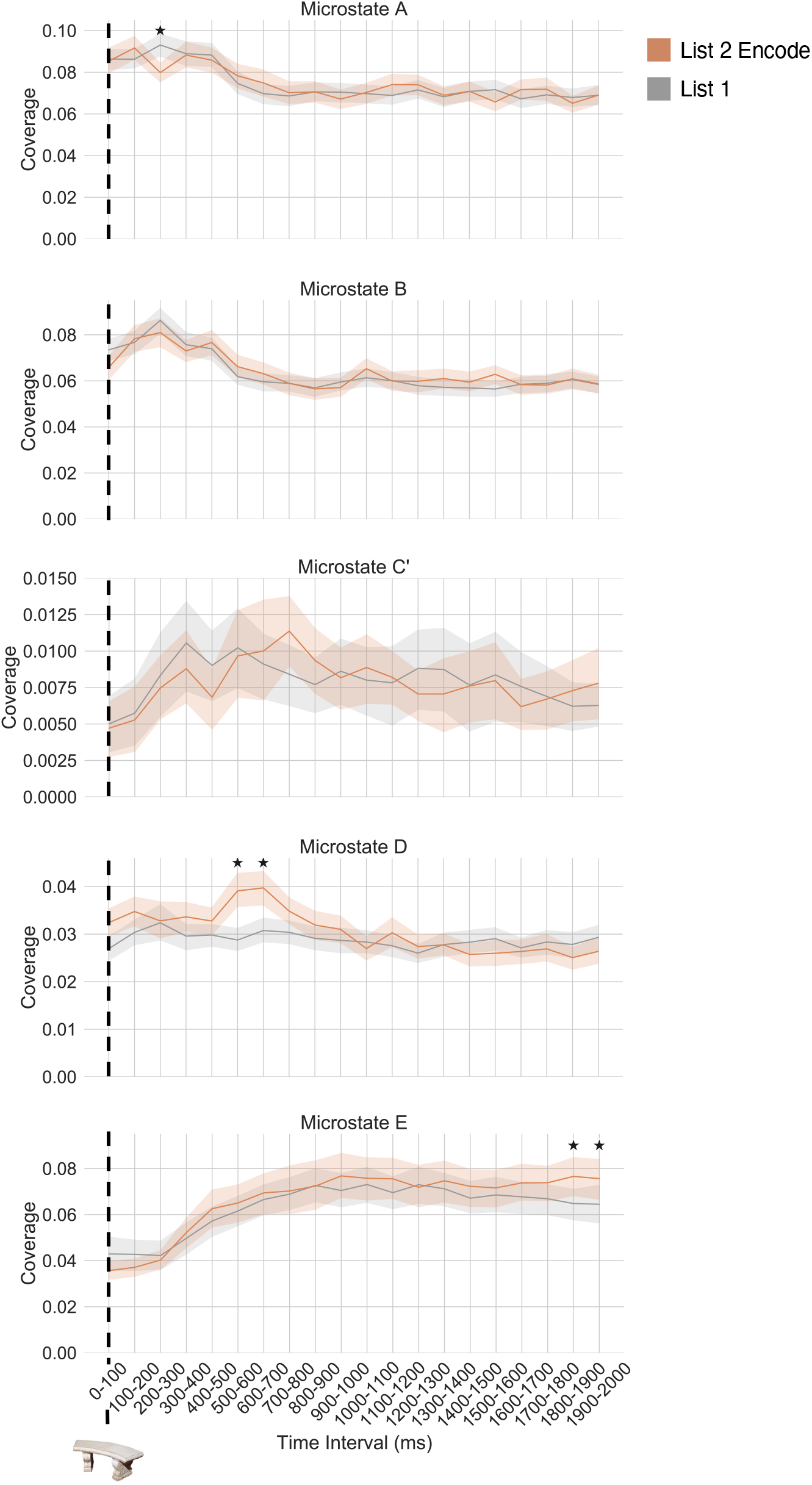
Temporal Dynamics of List 1 vs List 2 encode trials. We compared microstate coverage across List 2 encode (orange) and List 1 (grey) trials. Note that the y-axes differ across panels, as average microstate coverage (irrespective of condition) is variable. Stimulus onset at 0-100 ms is indicated by the vertical dashed line in each panel. The shaded area represents the standard error of the mean. * denotes time intervals for which the post-hoc *t* -test comparison between List 1 and List 2 encode trials survived FDR correction.

## Discussion

The goal of the present study was to investigate the spatiotemporal dynamics of memory states. Memory encoding and memory retrieval states can be dissociated by cortical network activity patterns (Long & Kuhl, 2019, 2021; Smith et al., 2022); however, whether memory states are supported by sustained activity in a specific network or by fluctuations across networks has remained an open question. By assessing the engagement of microstates, sustained global patterns of voltage scalp topographies (Michel & Koenig, 2018), we find that microstate E, a potential analog of the default mode network (DMN), specifically dissociates memory states and shows temporally sustained engagement during retrieval compared to encoding. Together, these results suggest that memory states, like other brain states, are supported by the maintenance of specific network configurations. These findings have implications for memory behavior and cognition more broadly, suggesting that hyper- or hypo-engagement of microstate E may lead to over or under engagement of a memory retrieval state.

We find that microstate E specifically dissociates memory encoding and memory retrieval states. Microstate E duration and coverage are significantly greater during retrieval compared to encoding. Thus, microstate E is more engaged and once active, stays engaged for longer during retrieve trials. Furthermore, we find that these dissociations are specific to microstate E through direct comparison with the other microstates identified in our resting state data. Prior work employing source localization suggests that microstate E may correspond to the DMN (Custo et al., 2017). Activity increases in the DMN during episodic memory retrieval (Kim, 2010), semantic processing (Binder et al., 2009), and internally directed attention (Buckner & DiNicola, 2019). In addition, activity patterns in the DMN also support retrieval-mediated prediction (Long, Lee, & Kuhl, 2016), accessing of schematic information (Preston & Eichenbaum, 2013; van Kesteren, Ruiter, Fernández, & Henson, 2012), and reliance on prior conceptual knowledge (González-García & He, 2021). Thus, our findings are highly consistent with the existing literature, as the retrieve condition in the current study may tap into any or all of these processes; subjects are instructed to retrieve a semantically associated stimulus encountered previously in the experiment. Which specific cognitive process microstate E engagement is supporting – or whether all of these processes constitute ‘internal mentation’ – is an important question for future work.

Microstate E engagement during retrieval is sustained throughout the stimulus interval. Aside from the first 200 ms following stimulus onset – when subjects are likely perceiving the stimulus in order to know which prior stimulus to retrieve – microstate E coverage is greater during retrieve compared to encode trials. Connectivity between the DMN and other cortical networks (e.g. the dorsal attention and frontoparietal control networks) is thought to support internal attention (Kam et al., 2019), working memory (Murphy et al., 2020), and search during free recall (Kragel & Polyn, 2013). The methodological approach for assessing microstates is winner-take-all such that only one microstate can be active at any given time sample. Thus, by definition it is not possible to demonstrate concurrent engagement of multiple microstates at the level of time samples. However, given the potentially multi-network involvement in memory retrieval, we might have anticipated changes in microstate coverage over time such that the dominant microstate would fluctuate throughout the stimulus interval. As an example, we might have expected to find transient changes in coverage between microstates B (putative visual network) and E, as subjects should use the visually presented stimulus as a cue to guide retrieval and could potentially alternate between externally directed attention to the current stimulus and internally directed attention to the retrieved stimulus. Although there may be conditions in which such fluctuations could be observed, in the current study with explicit instructions to retrieve a prior stimulus, we find that retrieval is characterized by sustained engagement of a single microstate. This finding has implications for future investigations into how memory states impact behavior. Hyper- or hypo-engagement of microstate E may result in too much or too little retrieval, respectively, which may negatively impact memory success.

Decreased microstate E engagement is a general property of an encoding state. Our mnemonic state task is designed such that on every trial with an explicit instruction, subjects *could* engage in either an encoding or retrieval state; the instructions are intended to bias subjects to one of these states. However, a consequence of this design is that signals during encode trials could reflect retrieval suppression rather than encoding per se. To demonstrate that less microstate E coverage is a general property of encoding, and not due to interference-mediated retrieval suppression, we measured microstate E coverage during trials in which retrieval was unlikely (if not impossible) and subjects’ goal was to encode the stimuli. We find that microstate E coverage is generally similar across both types of encoding trials, a result that is consistent with the interpretation that decreased microstate E engagement reflects a general encoding state, rather than top-down suppression of a retrieval state. However, the current study cannot address whether these effects extend beyond explicit or goal-directed memory states, as throughout the experiment, subjects had explicit goals to encode or retrieve. Microstate E and the DMN have been linked to spontaneous thought and mind wandering (Andrews-Hanna, Reidler, Sepulcre, Poulin, & Buckner, 2010; Bréchet et al., 2019; Higgins et al., 2021), suggesting that such signals may be a more general marker of the internal axis of attention (Chun, Golomb, & Turk-Browne, 2011). The engagement of microstates in incidental or automatic encoding and retrieval, as well as more broadly in the service of external vs. internal attention, is an important avenue for future research.

When subjects are instructed to encode, but have the potential to retrieve a prior stimulus, we find that encoding recruits microstate D. Specifically, microstate D coverage is greater when subjects are instructed to encode (and thereby not retrieve) in the 500-700 ms following stimulus onset. Microstate D may reflect the dorsal attention network (DAN; Britz et al., 2010; Custo et al., 2017), which supports the orienting and reorienting of attention based on top-down goals (Sestieri, Shulman, & Corbetta, 2017). One interpretation of the current findings is that elevated microstate D coverage reflects top-down control used to orient attention according to the task instructions, which is consistent with the occurrence of this dissociation early in the stimulus interval.

In summary, memory encoding and memory retrieval are differentiated by the sustained engagement of a putative default mode network microstate. The tendency to over or under engage this microstate may account for intra- and inter-individual variability in memory success. More broadly, sustained engagement of microstate E may impact other cognitive processes via its role in internally directed attention.

## Acknowledgments

Nicole Long is an iTHRIV Scholar. The iTHRIV Scholars Program is supported in part by the National Center for Advancing Translational Sciences of the National Institutes of Health under Award Numbers UL1TR003015 and KL2TR003016.

